# Comprehensive analysis of *Verticillium nonalfalfae in silico* secretome uncovers putative effector proteins expressed during hop invasion

**DOI:** 10.1101/237255

**Authors:** Kristina Marton, Marko Flajšman, Sebastjan Radišek, Katarina Košmelj, Jernej Jakše, Branka Javornik, Sabina Berne

## Abstract

**Background:** The vascular plant pathogen *Verticillium nonalfalfae* causes Verticillium wilt in several important crops. VnaSSP4.2 was recently discovered as a *V. nonalfalfae* virulence effector protein in the xylem sap of infected hop. Here, we expanded our search for candidate secreted effector proteins (CSEPs) in the *V. nonalfalfae* predicted secretome using a bioinformatic pipeline built on *V. nonalfalfae* genome data, RNA-Seq and proteomic studies of the interaction with hop.

**Results:** The secretome, rich in carbohydrate active enzymes, proteases, redox proteins and proteins involved in secondary metabolism, cellular processing and signaling, includes 263 CSEPs. Several homologs of known fungal effectors (LysM, NLPs, Hce2, Cerato-platanins, Cyanovirin-N lectins, hydrophobins and CFEM domain containing proteins) and avirulence determinants in the PHI database (Avr-Pita1 and MgSM1) were found. The majority of CSEPs were non-annotated and were narrowed down to 44 top priority candidates based on their likelihood of being effectors. These were examined by spatio-temporal gene expression profiling of infected hop. Among the highest *in planta* expressed CSEPs, five deletion mutants were tested in pathogenicity assays. A deletion mutant of *VnaUn.279*, a lethal pathotype specific gene with sequence similarity to SAM-dependent methyltransferase (LaeA), had lower infectivity and showed highly reduced virulence, but no changes in morphology, fungal growth or conidiation were observed.

**Conclusions:** Several putative secreted effector proteins that probably contribute to *V. nonalfalfae* colonization of hop were identified in this study. Among them, *LaeA* gene homolog was found to act as a potential novel virulence effector of *V. nonalfalfae*. The combined results will serve for future characterization of *V. nonalfalfae* effectors, which will advance our understanding of Verticillium wilt disease.

## 1 Background

Soil-born vascular plant pathogens, members of the *Verticillium* genus [1], cause Verticillium wilt in several economically important crops, including tomato, potato, cotton, hop, sunflower and woody perennials [2,3]. Studies of *Verticillium* – host interactions and disease processes, particular those caused by *V. dahliae,* have significantly contributed to the understanding of *Verticillium* spp. pathogenicity, although more research is needed for successful implementation of Verticillium wilt disease control [4–6].

Plant-colonizing fungi secrete a number of effectors, consisting among others of hydrolytic enzymes, toxic proteins and small molecules, to alter the host cell structure and function, thereby facilitating infection and/or triggering defense responses [7]. The assortment of effector molecules is complex and highly dynamic, reflecting the fungal pathogenic lifestyle [8] and leading to pathogen perpetuation and development of disease. Research into plant-pathogen interactions has significantly advanced, with an increasing number of sequenced microbial genomes, which have enabled computational prediction of effectors and subsequent functional and structural characterization of selected candidates. However, the prediction of fungal effectors, mainly small secreted proteins, which typically lack conserved sequence motifs and structural folds, is challenging and largely based on broad criteria, such as the presence of a secretion signal, no similarities with other protein domains, relatively small size, high cysteine content and species-specificity [8–10]. Using these features to mine predicted secretomes for candidate effectors has been valuable, but has not produced a one-size-fits-all solution [11]. Various approaches that combine several bioinformatics tools and also consider features such as diversifying selection, genome location and expression *in planta* [12–15], have had mixed outcomes. The EffectorP application has recently been presented as the first machine learning approach to predicting fungal effectors with over 80% sensitivity and specificity [16].

The genome sequences of five *Verticillium* species *(V. dahliae, V. alfalfae, V. tricorpus, V. longisporum* and *V. nonalfalfae)* and their strains have been published [17–22], providing a wealth of genomic information for various studies. Klosterman et al. [17] queried *V. dahliae* (strain VdLs.17) and *V. alfalfae* (strain VaMs.102) genomes for potential effectors and other secreted proteins based on subcellular localization and the presence of signal peptide. A similar number of secreted proteins was found in both genomes (780 and 759 for *V. dahliae* and *V. alfalfae,* respectively), and comparable to other fungi. These secretomes were further examined for effector candidates, obtaining 127 for *V. dahliae* and 119 for *V. alfalfae* proteins, based on the assumption that fungal effectors are small cysteine-rich proteins (SSCPs) with fewer than 400 amino acids and more than four cysteine residues. Siedl et al. [19] later re-examined the secretomes of two *V. dahliae* strains (VdLs.17 and JR2) and *V. alfalfae* (VaMs.102), and predicted a higher number of secreted proteins and smaller number of SSCPs, due to improved gene annotation and restricted criteria for SSCPs. Interestingly, in their comparison of highly pathogenic *Verticillium* species with saprophytic and weak pathogen *V. tricorpus,* a similar content of the secretome (cca 8.5%) in their respective proteomes was obtained. Orthologs of known effectors of *F. oxysporum, C. fulvum* or oomycete *Phytophtora infestans* were not found in *V. dahliae* and *V. alfalfae* genomes [17], except for *C. fulvum* LysM effector Ecp6 [23] (7 and 6 genes in *V. dahliae* and *V. alfalfae,* respectively) and *C. fulvum* virulence factor Ecp2 [24]. Several LysM effectors, widespread fungal proteins recognized by the LysM domain, have also been characterized as suppressors of PTI (PAMP-triggered immunity) through their chitin binding ability [25–28]. Reexamination of *Verticillium* LysM effectors has corroborated only three LysM effectors as core proteins in the genomes of *V. dahliae* strains with a function other than fungal pathogenicity, and one strain specific (VdLs.17) virulence associated LysM protein [18,27]. An increase in NLP (necrosis and ethylene-inducing protein (NEP-1)-like proteins) gene homologs has also been found among secreted proteins in *Verticillium* genomes (8 and 7 in *V. dahliae* and *V. alfalfae,* respectively) [17,19]. Zhou et al. [29] and Santhanam et al. [30] showed that only two of them displayed cytotoxic activity in tomato, cotton, *Arabidopsis* and *Nicotiana benthamiana,* while reduced virulence has been demonstrated for the deletion mutant of *VdNLP1* and *VdNLP2* in tomato and *Arabidopsis.*

Although numerous secreted proteins with unknown function or sequence similarity have been reported for *Verticillium* spp, only a handful have been characterized. Virulence effector Ave1 is a 134 aa long secreted protein with 4 conserved cysteines and an Expansin-like EG45 domain, discovered by comparative genomics in *V. dahliae* race 1 strains [31]. Ave1 recognition by tomato receptor-like protein Ve1 triggered immune signaling pathways leading to resistance to *V. dahliae* race 1 strains [32,33]. The other reported *V. dahliae* effector protein, PevD1, induced a hypersensitive response in tobacco [34,35] and triggered innate immunity in cotton plants, as demonstrated by upregulation of defense-related genes, metabolic substance deposition and cell wall modifications [36]. Zhang et al. [37] recently characterized a novel effector protein, VdSCP7, as a host nucleus targeting protein, which induced a plant immune response and altered the plant’s susceptibility to fungal and oomycete pathogens.

In addition to *V. dahliae* effectors, a small secreted protein, VnaSSP4.2, with an important role in fungal virulence, has been discovered in xylem sap during *V. nonalfalfae* infection of hops [38]. *V. nonalfalfae* is another pathogenic species of the genus *Verticillium*, although with a narrower host range than *V. dahlia* [1]. However, it causes Verticillium wilt and plant death in several important crops [4]. The occurrence of different virulent strains has been well documented in hop *(Humulus lupulus* L.), in which two pathotypes of *V. nonalfalfae* with different aggressiveness have been isolated, causing mild (fluctuating) and lethal (progressive) disease forms [39–43]. The disease is demonstrated by plant wilting, foliar chlorosis and necrosis, vascular browning and rapid plant withering and dieback in the lethal disease form [43]. Sporadic outbreaks of the *V. nonalfalfae* lethal pathotype in European hop gardens are of major concern, since there is no effective disease control except host resistance and strict phytosanitary measures. Despite the high economic losses caused by the lethal Verticillium wilt, the development of highly aggressive *V. nonalfalfae* pathotypes, as well as the genetics of hop resistance, remains enigmatic.

In the present study, a comprehensive biological database was generated using data from recently sequenced *V. nonalfalfae* genomes [22], transcriptomic and proteomic research of fungal growth on xylem stimulating medium [44] and RNA-seq studies of *V. nonalfalfae* – hop interactions [45]. A customized bioinformatics platform was used to set up a pipeline for prediction and characterization of the *V. nonalfalfae* secretome and select the best candidate secreted effector proteins (CSEPs) for functional studies. From a total of 263 CSEPs in the final dataset, the gene expression of the 44 highest ranking CSEPs was assessed by spatio-temporal RT-qPCR profiling of infected hop. Furthermore, deletion mutants of five selected CSEPs were analyzed in pathogenicity assays, with one of them exhibiting reduced virulence on hop plants. Our findings should assist further characterization of *V. nonalfalfae* effectors in an attempt to understand the molecular mechanisms of Verticillium wilt disease.

## 2 Results

### 2.1 The *V. nonalfalfae in silico* secretome is rich in carbohydrate-active enzymes, proteases and candidate secreted effector proteins (CSEPs)

The *V. nonalfalfae* genome comprises 9,269 predicted protein-encoding genes. Among these putative proteins, 944 are classically secreted proteins with signal peptide and no more than one transmembrane domain, representing 10.2% of the *V. nonalfalfae* predicted proteome (Figure 1 and Table S1). The accuracy of prediction was evaluated by comparing this dataset to a set of 91 unique sequences obtained by proteomic analysis (2D-DIGE) of *V. nonalfalfae* proteins secreted in xylem simulating medium [44], resulting in a 81.3% match (Table S1). Using TMHMM and Phobius [46] for transmembrane (TM) domain prediction, 801 proteins without a TM domain and 161 proteins harboring one TM domain were determined (Table S1). Based on subcellular localization predictions, 709 extracellular proteins (‘extr’ > 17) were acquired with WoLF PSORT [47], 450 proteins residing in the apoplast were determined with ApoplastP [48], while Localizer [49] identified 52 proteins harboring a chloroplast targeting signal and 12 proteins with a signal sequence for localization in mitochondria (Table S1).

**Figure 1.**
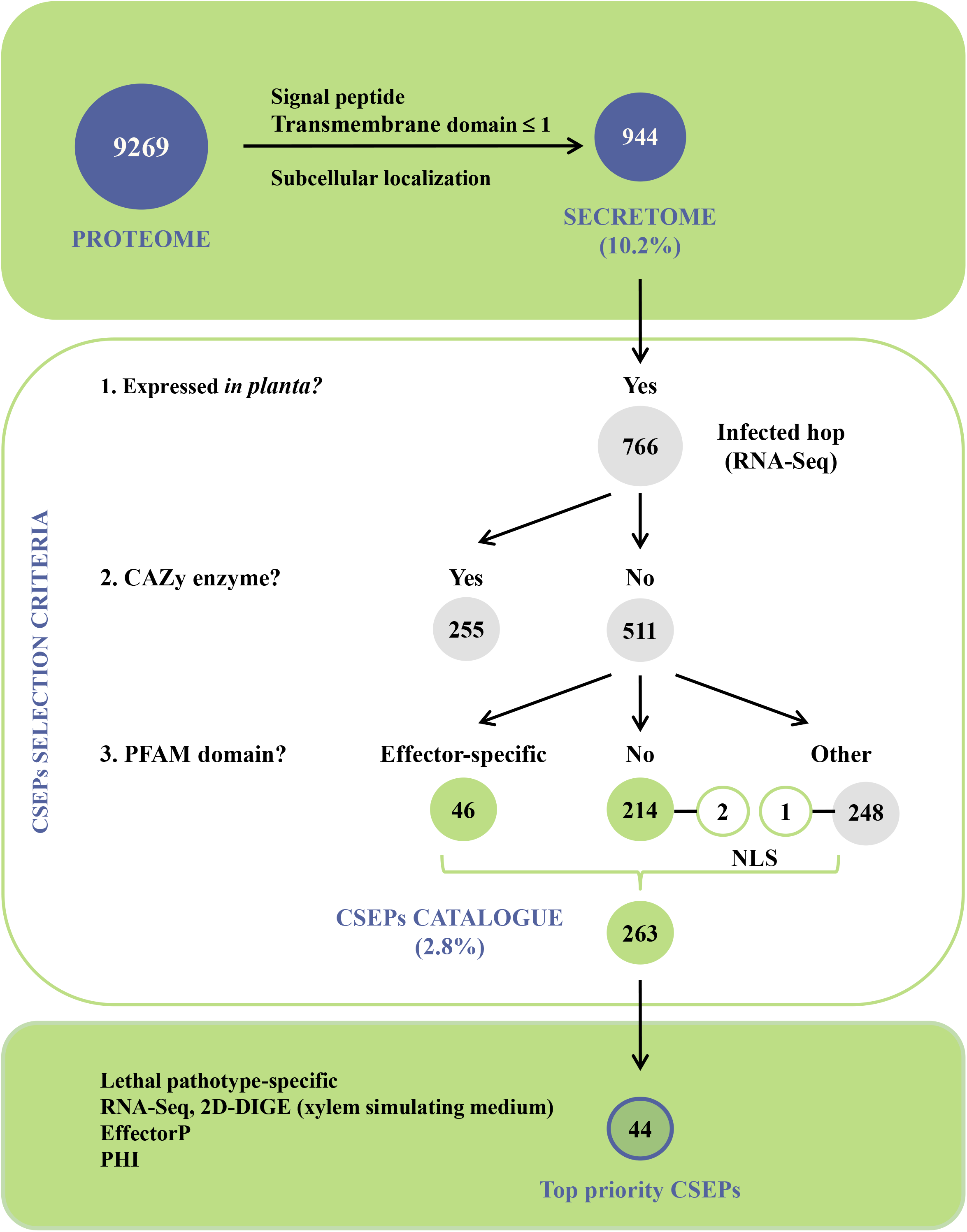
Bioinformatics pipeline for secretome prediction, identification and characterization of *V. nonalfalfae* candidate secreted effector proteins (CSEPs). *V. nonalfalfae* predicted proteome was first filtered based on signal peptide, transmembrane domains and subcellular localization to determine classically secreted proteins. This total predicted secretome was then refined to proteins expressed *in planta* and carbohydrate active enzymes were removed so that only proteins with effector-specific PFAM domains, NLS signal or no PFAM domains were retained in the final dataset of 263 CSEPs. These were narrowed down to 44 top-priority candidates based on the results of RNA-Seq and 2D-DIGE analyses, sequence similarity searching to known effectors in the PHI database and EffectorP prediction.

After similarity searches to known proteins in various databases, hypothetical functions were assigned to 727 (77%) putatively secreted *V. nonalfalfae* proteins. A superfamily annotation scheme [50] was used to classify the *V. nonalfalfae* secretome into seven functional groups (Figure 2). In the ‘Metabolism’ group, pectin lyase-like proteins and proteins with a cellulose-binding domain were over-represented within the ‘Polysaccharide metabolism and transport’ category, while (trans)glycosidases and six-hairpin glycosidases were predominant in the ‘Carbohydrate metabolism and transport’ category. Other abundant proteins in the ‘Metabolism’ group were reductases and heme-dependent peroxidases in the ‘Redox’ category, Concavalin A-like lectins/glucanases in the ‘Secondary metabolism’ category and ‘Transferases’. The major constituents of the ‘Intracellular processes’ group were ‘Proteases’ (especially acid proteases, Zn-dependent exopeptidases, subtilisin-like proteases and metalloproteases), cupredoxins in the ‘Ion metabolism and transport’ category and phospholipases in the ‘Phospholipid metabolism and transport’ category. Proteins involved in ‘Cell adhesion’ were over-represented within ‘Extracellular processes’, while most proteins in the ‘Information’ group classified into the ‘DNA replication/repair’ category. In the ‘General’ group, FAD-binding/transporter-associated domain-like proteins and proteins with a FAD/NAD(P)-binding domain were major constituents in the ‘Small molecule binding’ category. Analysis of the EuKaryotic Orthologous Groups (KOG) (Table S2) revealed a high number of proteins in the ‘Cellular processing and signaling’ category, associated with posttranslational modification, protein turnover, chaperones and signal transduction mechanisms, as well as cell wall/membrane/envelope biogenesis and intracellular trafficking, secretion, and vesicular transport, while the protein composition of the ‘Metabolism’ category mirrored that of the Superfamily.

**Figure 2.**
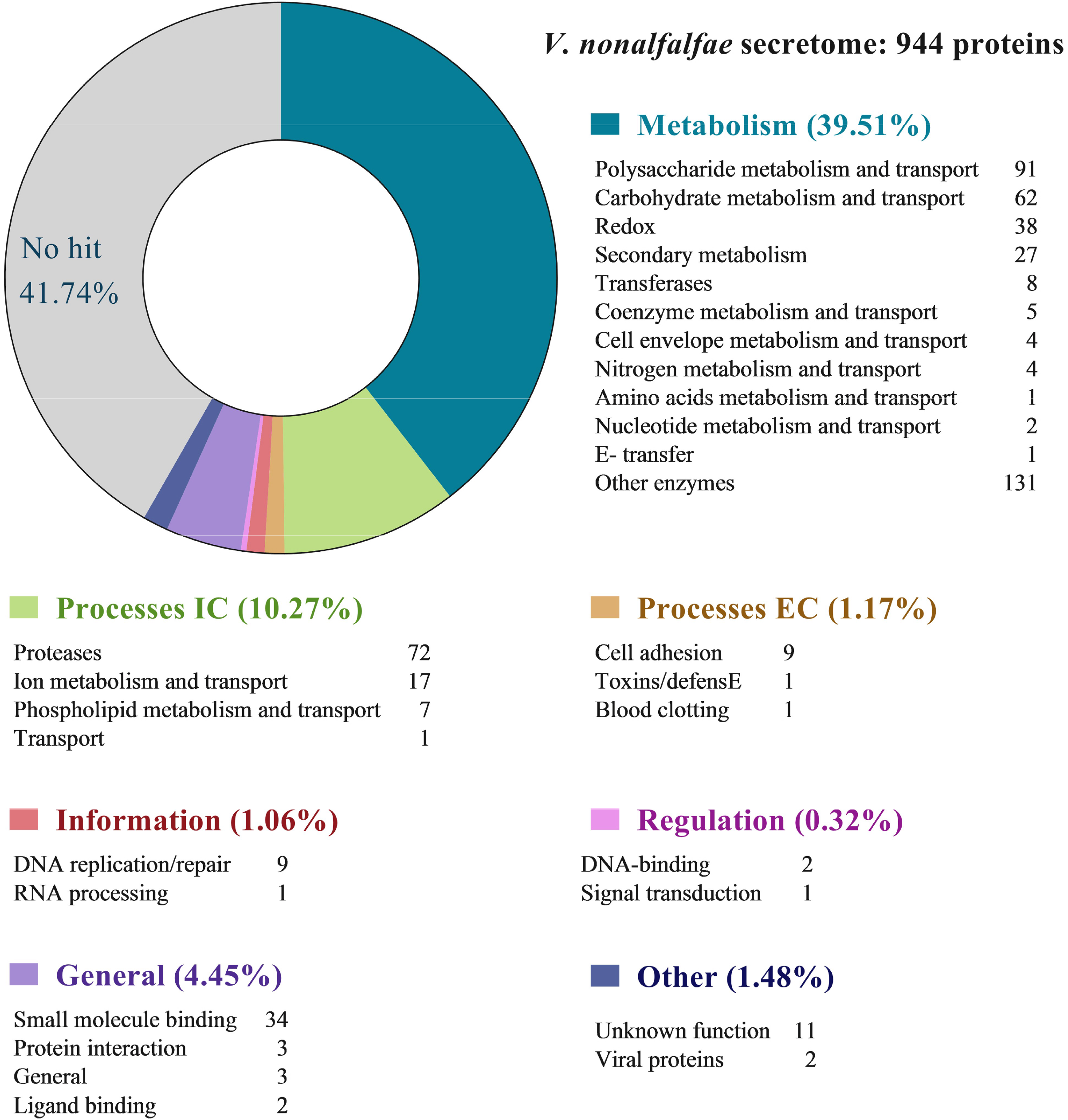
Classification of the *V. nonalfalfae* predicted secreted proteins into functional groups based on the Superfamily annotation scheme. Groups were classified after [50].

Since ‘Carbohydrate metabolic process’ and ‘Peptidase activity’ were overrepresented terms after Blast2GO analysis of the *V. nonalfalfae* secretome (Figure S1), they were investigated more thoroughly. Almost one third of the putative *V. nonalfalfae* secretome was CAZymes, of which 255 were expressed *in planta* and distributed as follows: glycoside hydrolases (129 GH), carbohydrate esterases (47 CE), redox enzymes that act in conjunction with CAZymes (49 AA; proteins with auxiliary activities), proteins with carbohydrate-binding modules (32 CBM), polysaccharide lyases (25 PL), and glycosyltransferases (4 GT). This repertoire of CAZymes was compared to other plant pathogenic *Verticillium* species (Figure 3A) and it was demonstrated that *V. nonalfalfae* had statistically more putative secreted CEs than *V. alfalfae*, in particular those involved in deacetylation of xylans and xylo-oligosaccharides. Moreover, the *V. nonalfalfae* secretome consisted of more putative secreted GHs than *V. dahliae* with major differences found in the GH3 group consisting primarily of stereochemistry-retaining β-glucosidases [51], the GH5 group of enzymes acting on β-linked oligo- and polysaccharides, and glycoconjugates [52], and the GH43 group of enzymes for the debranching and degradation of hemicellulose and pectin polymers [53]. In addition, the *V. nonalfalfae* secretome was enriched in putative secreted proteins with CBMs, when compared to *V. alfalfae, V. longisporum* and *V. dahlia.* This was largely due to cellulose-binding module CBM1 attached to various enzymes from families CE1, CE5, CE15, GH5, GH6, GH7, GH10, GH11, GH12, GH45, GH74, PL1, PL3 and AA9, and to some extent due to chitin-binding module CBM50 found in LysM effector proteins and subgroup C chitinases [54].

**Figure 3.**
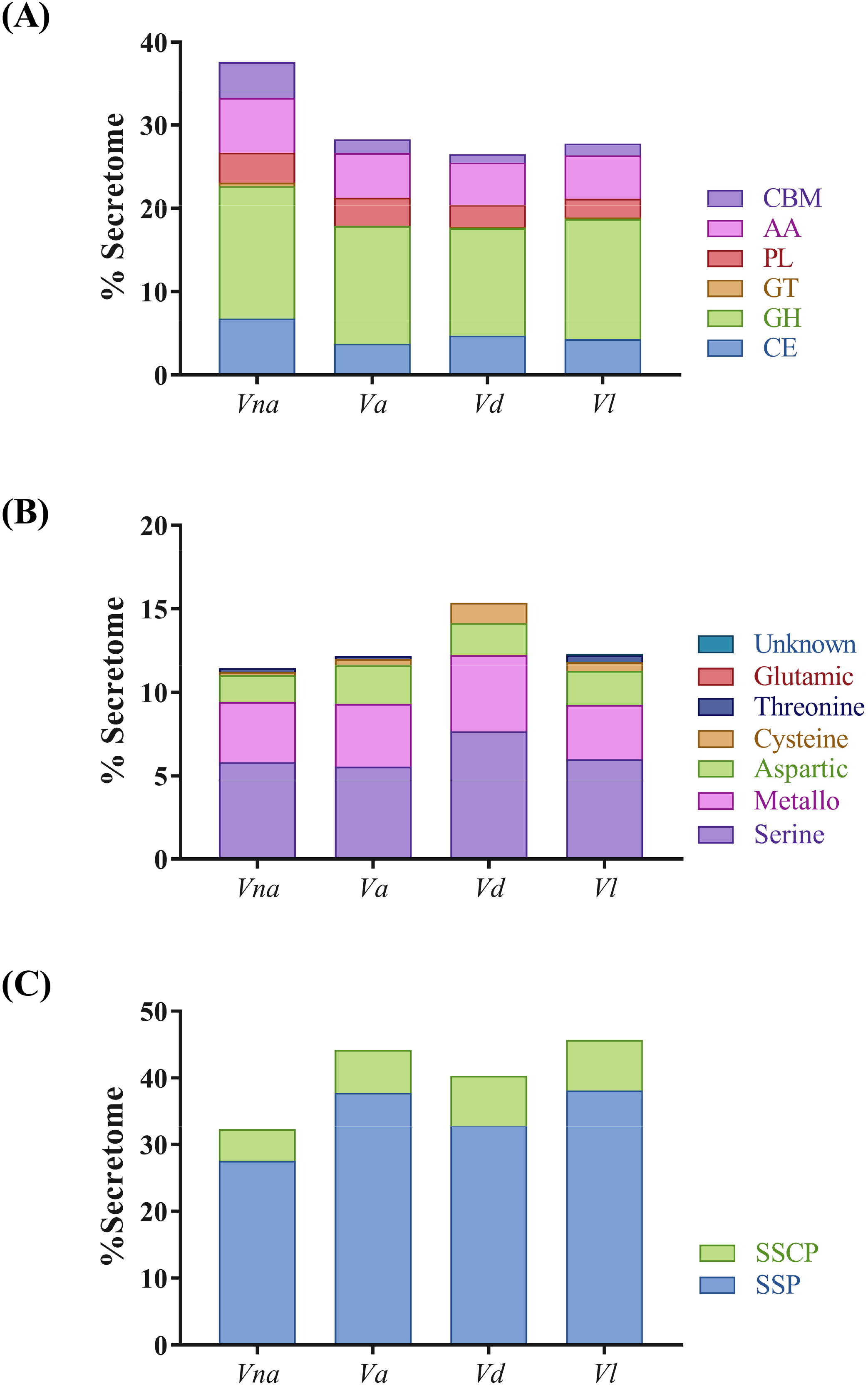
Relative abundance of carbohydrate-active enzymes (CAZymes), peptidases and small secreted proteins within secretomes of plant pathogenic *Verticillium* species. (A) Comparison of CAZymes from different classes is presented in percentages of predicted fungal secretome. CE, carbohydrate esterase; GH, glycoside hydrolase; GT, glycosyl transferase; PL, polysaccharide lyase; CBM, proteins with carbohydrate-binding modules; AA, proteins with auxiliary activities. (B) Comparison of various classes of peptidases depicted as percentage of predicted fungal secretome. (C) Relative abundance of small secreted proteins and small secreted cysteine rich proteins in predicted fungal secretomes. SSPs, small secreted proteins with less than 300 aa; SSCPs, small secreted proteins with more than 5% cysteine content and at least 4 Cys residues [55]; *Vna, V. nonalfalfae*; *Va*, *V. alfalfae*; *Vd*, *V. dahliae*; *Vl*, *V. longisporum*.

Similarity searching of *V. nonalfalfae* putative secreted proteins against peptidases in the *MEROPS* database revealed 12 *in planta* expressed aspartic peptidases, 2 cysteine peptidases, 27 metallopeptidases, 44 serine peptidases and 1 threonine peptidase. The highest representation of *V. nonalfalfae* putative secreted peptidases was in the M14A (carboxypeptidase A1), S08A (subtilisin Carlsberg) and A01A (pepsin A) subfamilies (Table S3). Comparison of putative secreted peptidases between plant pathogenic *Verticillium* species (Figure 3B) revealed a similar distribution of peptidases among *V. nonalfalfae, V. alfalfae* and *V. longisporum,* while *V. dahliae* had a statistically different distribution of metallopeptidases, cysteine and serine peptidases. Other enzymatic activities of putatively secreted *V. nonalfalfae* proteins according to the KEGG analysis can be found in Table S4.

Querying *Verticillium in silico* secretomes for small secreted proteins (SSPs) of less than 300 aa and small secreted cysteine rich (SSCPs) proteins with more than 5% cysteine content and at least 4 Cys residues [55] showed 5-10% lower abundance of SSPs and 2-3% fewer SSCPs in the *V. nonalfalfae* secretome than in the secretomes of other plant pathogenic *Verticillium* species (Figure 3C). Since small secreted proteins are the least characterized portion of fungal secretomes and many have been shown to act as effectors, our secretome analysis further focused on filtering secreted proteins for expression *in planta* to identify CSEPs relevant to *V. nonalfalfae* infection of hop. Genome-wide transcriptome analysis of the *V. nonalfalfae* interaction with hop [45] revealed that 766 (81%) transcripts in the *V. nonalfalfae in silico* secretome were expressed in infected hop samples. They showed distinct expression patterns related to different stages of infection (6, 12, 18 and 30 dpi), hop cultivar (susceptible ‘Celeia’ or resistant ‘Wye Target’) and plant tissue (roots or shoots) (Figure S2). From this dataset, all CAZymes except CBMs were omitted from further analysis, resulting in 529 putatively secreted *in planta* expressed proteins (Figure 1), of which 308 had sequence similarity to PFAM domains (Table S5). These included, among others, effector-specific PFAM domains, such as LysM effectors [27], Necrosis inducing proteins (NPP1) [56], Hce2 (Homologs of *Cladosporium fulvum* Ecp2 effector) effector proteins [57], Cerato-platanins [58], Cyanovirin-N lectins [59], hydrophobins [60] and CFEM (Common in Fungal Extracellular Membranes) domain containing proteins [61]. Our final dataset of CSEPs comprised a total of 263 proteins without functional PFAM domains, and proteins bearing known effector-specific PFAM domains, representing 2.8% of the putative proteome. Among them, we determined also 3 CSEPs with a nuclear localization signal (NLS), implying their activity in the plant nucleus, 3 CSEPs specific to the lethal strain of *V. nonalfalfae* and 69 probable effector proteins (Table S6) as predicted by EffectorP [16]. Similarity searching of CSEPs to experimentally verified pathogenicity, virulence and effector genes from fungal, oomycete and bacterial pathogens in the Pathogen-host interaction (PHI) database [62] revealed proteins matching AVR effectors (5 hits) and known effector proteins displaying reduced virulence (11 hits) or unaffected pathogenicity (4 hits) (Table S6).

### 2.2 *V. nonalfalfae* CSEPs display distinct gene expression profiles during infection of hop

Establishing successful colonization of a host plant requires effective and timely delivery of the fungal pathogen’s effectors. Using quantitative real-time PCR, the expression of the 44 top-priority CSEPs selected according to their likelihood of being effectors (see Methods for selection criteria), was investigated in root and shoot samples of Verticillium wilt susceptible (‘Celeia’) and resistant (‘Wye Target’) hop at 6, 12 and 18 days after inoculation with *V. nonalfalfae*. In a preliminary experiment, the average expression of the selected CSEPs in pooled root samples at different time points was examined (Table S7). The three highest expressed CSEPs that were also selected by Effector P prediction, and two lethal pathotype specific CSEPs, were then profiled using biological replicates (Figure 4). We included the *VnaSSP4.2* gene, encoding a small secreted protein, in the gene expression analysis as a positive control for virulence-associated *V. nonalfalfae* effector [38].

**Figure 4.**
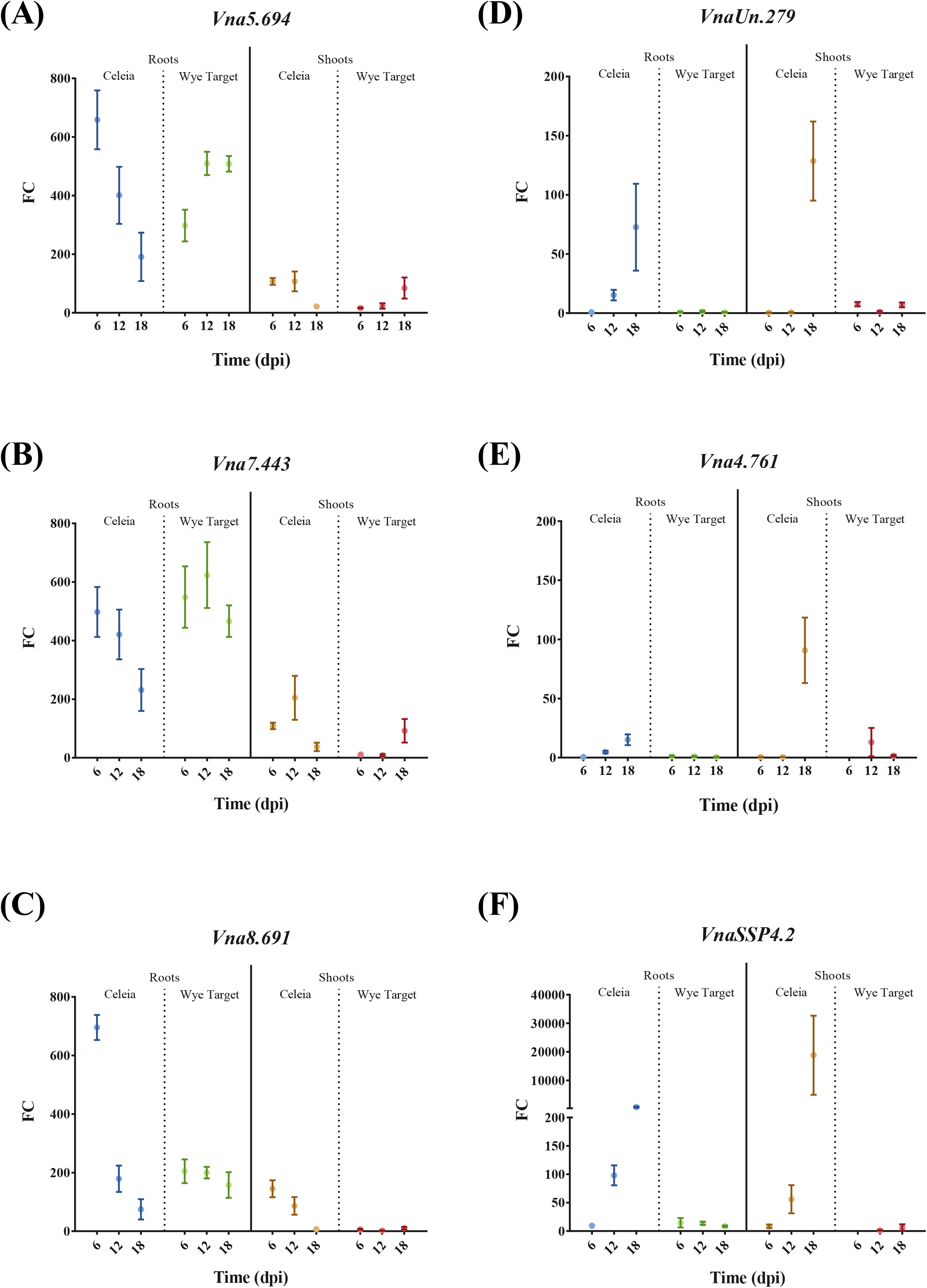
Gene expression profiles of selected *V. nonalfalfae* CSEPs in the roots and shoots of infected hop. FC, fold change in gene expression was determined by quantitative real-time PCR using topoisomerase and splicing factor 3a2 as endogenous controls and fungal samples grown on ½ CD medium as a reference. Means ± SEM (n=5) are presented. Statistical significance was determined with the t-test using the Holm-Sidak approach, with *α* = 5%. ‘Celeia’, *Verticillium* wilt susceptible hop; ‘Wye Target’, *Verticillium* wilt resistant hop; dpi, days post inoculation

The expression levels of genes *Vna5.694, Vna7.443* and *Vna8.691* were greater in the roots than in the shoots of both hop varieties. Gene expression of Vna5.694 (Figure 4A), encoding a small (81 aa) secreted cysteine-rich protein of unknown function and displaying the highest similarity to *V. longisporum* CRK15920 protein, decreased with time in the roots of susceptible hop, while its expression in the resistant hop increased. A similar trend of expression was also observed in the shoots of both hop varieties. Gene expression of *Vna7.443* (Figure 4B), which produces a secreted protein (276 aa without cysteines) with the highest similarity to *V. longisporum* CRJ82870 protein, was comparable to *Vna5.694*; it decreased in the roots of susceptible hop and peaked at 12 dpi in the roots of resistant hop. A peak of expression at 12 dpi was also observed in the shoots of susceptible hop, while its expression increased with time in the shoots of resistant hop. Expression of the *Vna8.691* gene (Figure 4C), coding for a small secreted protein (95 aa without cysteines) of unknown function with the highest similarity to *V. longisporum* CRK10461 protein, was highest in the roots of susceptible hop at 6 dpi and then decreased with time of infection. The same trend was also observed in the shoots of susceptible hop. Expression of *Vna8.691* in the roots of resistant plants was constant, and around 200-fold higher than that in ½ CD medium, whereas no expression was detected in the shoots. Interestingly, two lethal pathotype specific genes *VnaUn.279* (Figure 4D), encoding a small secreted protein (92 aa without cysteines) with the highest similarity to *V. dahliae* VdLs.17 EGY23483 protein, and *Vna4.761* (Figure 4E), encoding a 186 aa protein with 8 cysteines and the highest similarity to *V. longisporum* CRK16219 protein, had similar gene expression patterns as the virulence-associated *V. nonalfalfae* effector *VnaSSP4.2* (Figure 4F). They were expressed only in the roots and shoots of susceptible hop (with expression levels increasing with time of infection) and barely detected in the resistant plants.

### 2.3 Identification of novel virulence effector of *V. nonalfalfae*

Since all five selected CSEPs were specifically expressed during plant colonization, a reverse genetics approach was used to test their contribution to the virulence of *V. nonalfalfae* in hop. Knockout mutants of *VnaUn.279* displayed only minor wilting symptoms in the susceptible hop (Figure 5), while vegetative growth, fungal morphology and sporulation were not affected. Analysis of the relative area under the disease progress curve (rAUDPC) [63] indicated that four independent *VnaUn.279* knockout mutants displayed statistically significantly lower values of rAUDPC than the wild type fungus (Figure 6A). To understand the progress of disease in time, statistical modelling was undertaken. For illustrative purposes only, disease severity index (DSI) values [42] for *AVnaUn.279* and wild type fungus were modelled by logistic growth model (Figure 6B). The variability of disease progression in individual hop plants is probably due to the specific nature of *Verticillium* colonization, in which only a few attached hyphae randomly penetrate the root intercellularly [64]. As demonstrated by the inflection point of the *ΔVnaUn.279* logistic curve, development of disease symptoms was delayed for 10 days compared to the wild type fungus. Based on the asymptote values, the wilting symptoms were considerably less severe in the mutant than in the wild type fungus. Additionally, fungal biomass assessment with qPCR revealed that 32% of plants were infected with *V. nonalfalfae Δ VnaUn.279* mutants compared to at least 80% for the wild type fungus. These results indicate that deletion of *VnaUn.279* not only severely reduced *V. nonafalfae* virulence but also significantly affected the fungal infectivity via a yet unknown mechanism.

**Figure 5.**
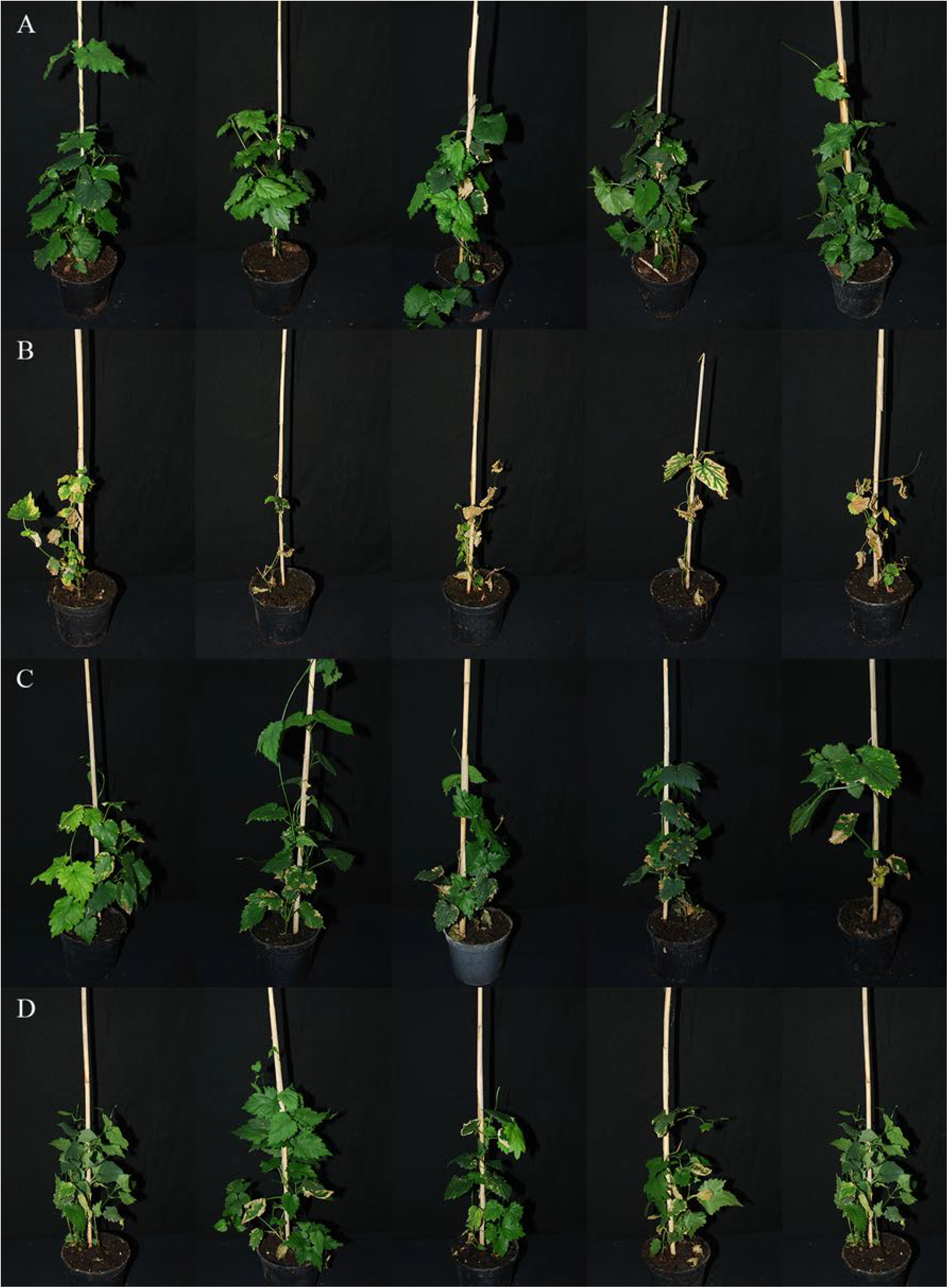
Pathogenicity testing in susceptible hop ‘Celeia’. Pictures were taken 31 days post inoculation of hop roots with *V. nonalfalfae* conidia suspension. (A) mock-inoculated plants; (B) plants inoculated with wild type *V. nonalfalfae* T2 strain; (C and D), plants inoculated with *V. nonalfalfae ΔVnaUn.279* mutant strains (two replicates).

**Figure 6.**
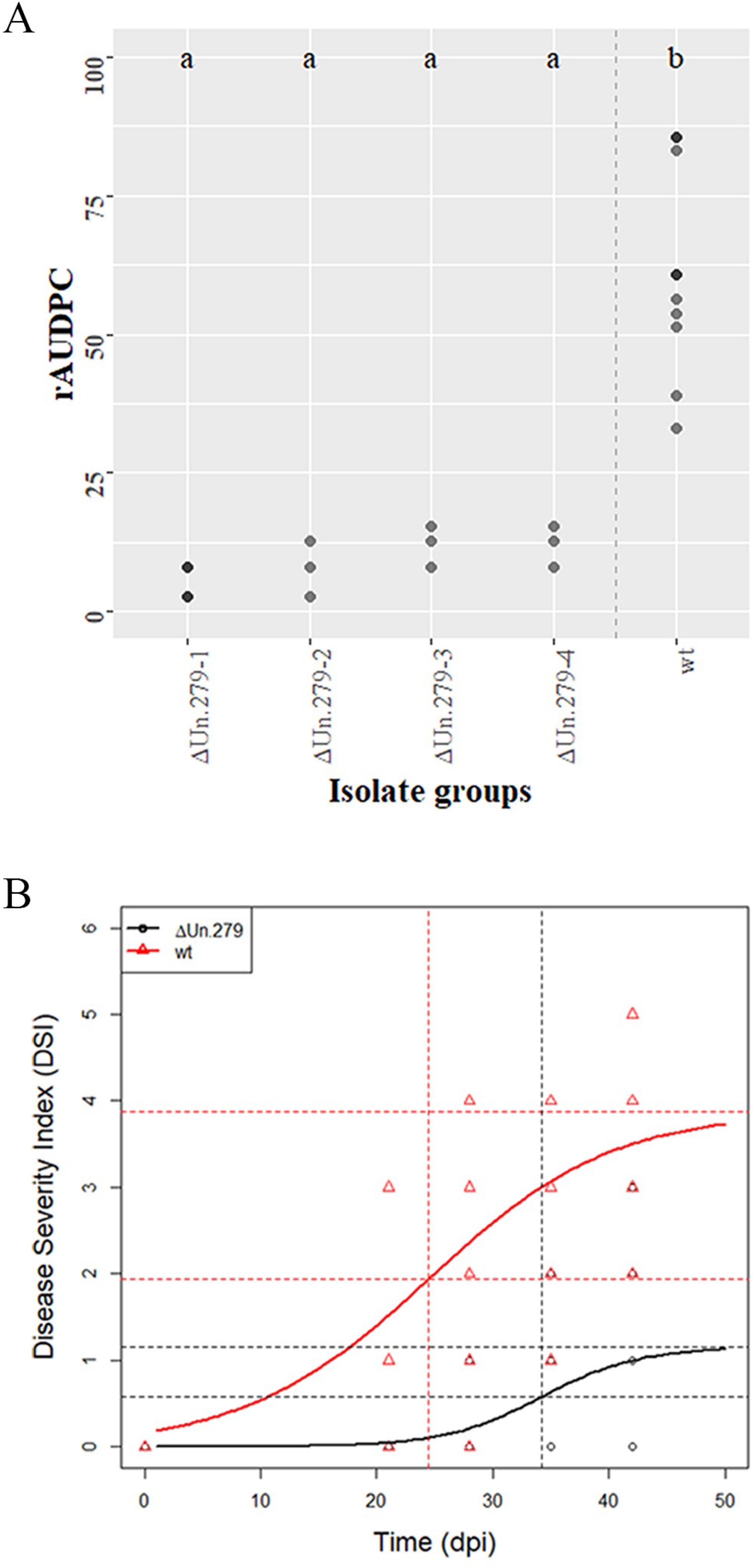
*V. nonalfalfae* knockout mutant Δ*VnaUn.279* showed severely reduced virulence. (A) Relative area under the disease progress curve (rAUDPC) is presented for knockout mutant *ΔVnaUn.279* and for wild type (wt) *V. nonalfalfae.* Darker dots depict double values. A Kruskal-Wallis test followed by multiple comparison test resulted in two groups: *a* for Δ*VnaUn.279* and *b* for wild type. (B) Disease severity index (DSI) values were fitted by a simple logistic growth model for mutant Δ*VnaUn.279* and for wild type *V. nonalfalfae* hop isolates. The upper horizontal line is the asymptote (black for Δ*VnaUn.279,* red for wt); the vertical lines show the inflection points (black for Δ *VnaUn.279,* red for wt) at which the predicted DSI is one half of the asymptote.

The other tested *V. nonafalfae* knockout mutants *(Vna5.694, Vna7.443* and *Vna8.691* in Figure S3) showed unaffected pathogenicity and no statistical differences in rAUDPC values relative to the wild type. Deletion of *Vna4.761* was not achieved, due to its functional redundancy, since two additional copies with over 95% sequence identity have been found after Blastn search against *V. nonalfalfae* reference genome at two different genomic locations.

## 3 Discussion

Fungal pathogens have evolved diverse strategies to interact with host plants and secrete various effector molecules to overcome plant defense mechanisms. A recently published genome of xylem-invading Sordariomycete fungus *Verticillium nonalfalfae* [22], a transcriptome study of infected hop [45] and obtained proteomic data of fungal growth on xylem simulating medium [44] have provided an opportunity to screen for proteins that may contribute to fungal virulence in hop. In the current study, a customized bioinformatics pipeline was designed to predict the classical *V. nonalfalfae* secretome and then to refine the secretome based on experimental data to identify candidate secreted effector proteins (CSEPs).

The relative secretome size of *V. nonalfalfae* (10.2%) conforms to the nutritional lifestyle of plant pathogens with larger secretome sizes (from 2.9% to 12.1% of the proteomes, with an average of 7.4%) and fits in the phylogenetic context with other Pezizomycotina (from 3.7% to 12.1% of the proteomes, with an average of 7.3%) [55]. The majority of proteins (69.5%) in the *V. nonalfalfae* secretome were less than 500 aa residues. In contrast to some plant pathogenic Pezizomycotina, which had remarkable 10-15% enrichment of proteins of up to 100 aa residues [55], only 1.8% of such proteins were found in the *V. nonalfalfae* secretome.

The composition of the *V. nonalfalfae* predicted secretome, rich in carbohydrate active enzymes (33%), proteases (11%), lipases/cutinases (4.6%) and oxidoreductases (4%), reflected its nutritional lifestyle as a hemibiotroph and plant vascular pathogen. Hemibiotrophic fungi undergo two phases during the infection process; an initial biotrophic phase, with characteristic expression of small secreted proteins without functional annotation (SSPs), is followed by a necrotrophic stage, which is generally associated with the expression of plant cell wall-degrading enzymes (CWDEs) [8]. Similar to *V. dahliae* and *V. alfalfae* genomes [65], the *V. nonalfalfae* genome encodes more CWDEs per number of secreted proteins than other plant pathogenic fungi [8]. Pectinases, xylanases, cellulases, glucanases, proteases, cutinases and lipases are major classes of CWDEs [66] and play important roles during plant colonization. They may facilitate penetration of the plant roots to reach the xylem vessels, degrade pectin gels and tyloses, formed in response to infection, to spread inside vessels, breakdown insoluble wall polymers to acquire nutrients and contribute to the release of survival structures from dead plant material [65]. In addition to contributing to virulence [67], some CWDEs are recognized as pathogen-associated molecular patterns (PAMPs), which provoke PAMP-triggered immunity [68,69]. On the other hand, *V. dahliae* carbohydrate-binding module family 1 domain-containing proteins may suppress glycoside hydrolase 12 protein-triggered immunity in plants [70]. As in the case of *V. dahliae* [65], the *V. nonalfalfae* predicted proteome contains numerous glycoside hydrolases and polysaharide lyases, most of which are secreted [44]; however, only five glycosyl transferases were determined in the predicted secretome. In contrast to *V. dahliae,* there was almost double the number of carbohydrate esterases in the *V. nonalfalfae* proteome and over half were predicted to be secreted. Other abundant proteins in the *V. nonalfalfae* predicted secretome were acid proteases, subtilisin-like proteases and zinc-dependent metalloproteases. These enzymes probably participate in amino acid acquisition, manipulation of host defenses by degradation of pathogenesis-related proteins, including plant chitinases, and act as virulence factors or as elicitors of defense responses [71,72]. A significant number of lipases, phospholipases and cutinases were determined in the *V. nonalfalfae* predicted secretome and identified in a previous proteomic study [44]. In addition to supplying energy for pathogen growth, lipid hydrolysis is crucial for the production of certain signaling molecules, such as oxylipins, which manipulate the host lipid metabolism and alter plant defense responses [73]. The role of cutinases in pathogenicity is controversial and has been associated with the dissolution of the plant cuticle during penetration, suppression of callose formation, spore attachment and surface signaling [74,75]. Another group of abundant enzymes in *V. nonalfalfae* predicted and experimentally determined secretomes were oxidoreductases, in particular FAD-dependent oxidoreductases and GMC oxidoreductases. Oxidoreductases are probably secreted for protection against host-produced reactive oxygen species, such as the generation of H2O2, which was detected after infection with *V. dahliae* in cotton roots [76] and in tomato plants [77]. On the other hand, fungal pathogens can actively contribute to the ROS level in host plants [78]. In *V. dahliae,* NADPH oxidase complex (Nox), composed of the catalytic subunit VdNoxB and tetraspanin VdPls1, is responsible for the production of ROS and the formation of penetration peg within the hyphopodium [64]. Moreover, VdNoxB regulates the cytoskeletal organization of the VdSep5-septin ring that separates the hyphopodium from invasive hyphae and forms a specialized fungus-host penetration interface, where small secreted proteins preferentially accumulate [79].

Since small secreted proteins (SSPs) are the least characterized fungal secreted proteins and some have been reported as effectors, we particularly focused on this group of proteins. Various criteria for the determination of SSPs have been reported [12,80,81] but, for comparison purposes, we adopted a definition [55] that considers SSPs proteins with a mature length of ≤ 300 aa residues and proteins with a relative cysteine content of ≥ 5%, as well as ≥ 4 cysteine residues, to be small secreted cysteine-rich proteins (SSCP). According to these criteria, the *V. nonalfalfae* predicted secretome contains 310 SSPs (32.8% of the predicted secretome) and 46 (4.9% of the secretome) of those belong to SSCPs. These numbers are lower than the average contents of SSPs (49%) and SSCPs (6.7%) determined in fungi of class 2 secretome size (500-1100 secreted proteins) and average contents of SSPs (47%) and SSCPs (7.5%) in the Pezizomycotina group [55]. In a recent study, SSPs with a mature length of ≤ 300 aa residues were identified in 136 fungal species and compared in terms of taxa and lifestyles [82]. On average, hemibiotrophs and necrotrophs had higher proportions of secreted enzymes, while biotrophs, symbionts and certain hemibiotrophs had the most abundant SSPs. Furthermore, higher numbers of species-specific SSPs (over 100) were associated with biotrophs and symbionts than necrotrophs and saprotrophs (around 30), suggesting that these effectors coevolved with their hosts, while the range was widest for hemibiotrophs. However, no species-specific SSPs have been discovered in *V. nonalfalfae,* while 13 and 19 have been reported in *V. albo-atrum* and *V. dahliae,* respectively [82].

Further analysis of the refined *V. nonalfalfae* secretome, comprised of 263 CSEPs, focused in particular on homologs of known effectors from other plant pathogens. Searching for LysM effectors using CAZy module (CBM50) and PFAM domain PF01476 revealed that the *V. nonalfalfae* secretome contains four *in planta* expressed proteins with 2-6 LysM domains, of which Vna2.979 is an ortholog of VDAG_00902 (Table S6), a core LysM effector of *V. dahliae* [27]. The necrosis- and ethylene-inducing-like proteins (NLPs) are a group of widespread conserved effectors that can trigger immune responses and cell death [56]. Similar to other *Verticillium* spp. [19,65], NLP genes are expanded in the *V. nonalfalfae* genome, with seven genes orthologous to *V. dahliae* NLP 1-9 and having no ortholog to VdNLP6. All five NLPs with homologs in the PHI database (Table S6) were expressed in hop; the most abundant was Vna7.239 (VdNLP9), in particular in the roots of susceptible hop (Figure S2), suggesting some role in the plant colonization process. Four fungal hydrophobins, characterized by high levels of hydrophobicity and the presence of eight conserved cysteine residues [60], were found in the refined *V. nonalfalfae* secretome and had homologs in the PHI database (Table S6). Although all were expressed in hop, only expression of Vna7.87 was abundant and root-specific. Interestingly, a role in the development of microsclerotia [83] was demonstrated for type II hydrophobin VDH1 from *V. dahliae,* but it was not required for pathogenicity. Further mining of CSEPs for known effectors revealed that *in planta* expressed Vna2.8 and Vna3.54 are homologs of AVR-Pita1, a neutral zinc metalloprotease from *Magnaporthe oryzae* [84], and Vna1.1274 was similar to MgSM1, a putative small protein of the Cerato-platanin family [85]. Three *V. nonalfalfae* CSEPs, Vna10.263 and Vna5.719 with high and ubiquitous expression in hop, and Vna9.246 specifically expressed only in susceptible hop, had similarity to *Candida albicans* RBT4, secreted pathogenesis-related proteins [86], while one CSEP was an ortholog of urea amidolyase (DUR1,2), which enables utilization of urea as a sole nitrogen source [87]. Highly expressed secreted protein Vna4.130 had similarity to EMP1, extracellular matrix protein 1 from *Magnaporthe grisea,* which was required for appressorium formation and pathogenicity [88]. Vna7.617, which was putatively secreted and abundantly expressed in susceptible hop, had an ortholog in *Fusarium oxysporum* membrane mucin Msb2, which regulates invasive growth and plant infection upstream of Fmk1 MAPK [89].

The remaining *V. nonalfalfae* CSEPs (92%) were hypothetical, predicted and conserved hypothetical proteins with no functional annotation and their temporal gene expression patterns in susceptible and resistant hop were explored to provide some clues to their function (Table S7 and Figure S2). In addition to the already reported effector VnaSSP4.2 [38], several other CSEPs with distinct expression patterns and high levels of expression were found. Among five CSEPs selected for gene functional analysis using a reverse genetics approach, four were predicted as effectors by EffectorP [16] and one had an ortholog in the PHI database, displaying sequence similarity to fungal effector LaeA, a regulator of secondary metabolism [90] and morphogenetic fungal virulence factor [91]. After comparing the pathogenicity of wild type fungus to CSEPs knockout mutants (Figure 6 and Figure S3), we discovered that the later CSEP, encoded by lethal pathotype specific gene *VnaUn.279,* is a novel virulence factor of *V. nonalfalfae. Δ VnaUn.279* mutants had diminished infectivity and exhibited severely reduced virulence in hop. Reduced virulence was also reported for a number of *LaeA* deletion mutants from human pathogen *A. fumigatus* [91], plant pathogenic fungi, including *A. flavus, C. heterostrophus* and several *Fusarium* species [92], as well as entomopathogenic fungus *Beauveria bassiana* [93]. Alltogether, these findings justify further investigation of the biological role of VnaUn.279 in *V. nonalfalfae* pathogenicity.

Despite the other selected CSEPs mutants not displaying any virulence associated phenotype, based on their expression profiles, they probably participate in other physiological processes during *V. nonalfalfae* infection of hop. Additionally, certain CSEPs may be recognized (and subsequently termed Avr effectors) by plant resistance proteins (R proteins), which are intracellular nucleotide-binding leucine rich repeat (NLR) receptors, via direct (receptor-mediated binding) or indirect (accessory protein-mediated) interactions, resulting in effector triggered immunity (ETI) [94,95]. To support this hypothesis, further testing of CSEPs mutants in the resistant hop cultivar is required and could result in identification of corresponding hop resistance proteins. These may then be exploited in Verticillium wilt control by introducing new genetic resistance traits into hop breeding, as already successfully implemented in certain other crops [96].

## 4 Conclusions

After comprehensively investigating the predicted *V. nonalfalfae* secretome using a diverse bioinformatics approaches and integrating multiple lines of evidence (genomics, transcriptomics and proteomics), several candidate secreted effector proteins were identified among protein-encoding genes. These are of high interest to scientists working on Verticillium wilt and, more generally, on pathogen effectors. Since the majority were non-annotated protein sequences, two strategies were adopted to gather clues about their function. With spatio-temporal gene expression profiling, we identified those candidate effectors that have important roles during *V. nonalfalfae* colonization of hop, while pathogenicity assays with effector knockout mutants revealed the candidates that contribute to fungal pathogenicity in hop. In conclusion, a new virulence effector of *V. nonalfalfae*, encoded by lethal-pathotype specific gene *VnaUn.279*, was identified and will be subject to future functional and structural studies.

## 5 Material and Methods

### 5.1 Microbial strains and cultivation

Sordariomycete fungus *Verticillium nonalfalfae* [1] was obtained from the Slovenian Institute of Hop Research and Brewing fungal collection. Two isolates from infected hop were used, differing in aggressiveness: lethal pathotype (isolate T2) and mild pathotype (isolate Rec) [97]. Fungal mycelium was cultured at room temperature on a half concentration of Czapek Dox broth (½ CD), supplemented with 1 g/L malt extract, 1 g/L peptone (all from Duchefa, The Netherlands) and 1 g/L yeast extract (Sigma Life Science, USA). For solid media, 15 g/L agar (Duchefa, The Netherlands) was added to ½ CD with supplements. Alternatively, potato dextrose agar (PDA; Biolife Italiana Srl, Italy) or Xylem simulating medium (XSM; [98]) was used.

*Escherichia coli* DH5α, used for amplification of vector constructs, was cultivated in LB medium with 50 mg/L kanamycin (Duchefa, The Netherlands) at 37°C.

*Agrobacterium tumefaciens* (LBA4404) transformation was performed in YM medium [99] containing 100 mg/L streptomycin and 50 mg/L kanamycin (both from Duchefa, The Netherlands) at 30°C. The co-cultivation of transformed *A. tumefaciens* and *V. nonalfalfae* was carried out on IMAS plates [99] at room temperature.

### 5.2 Functional annotation of *V. nonalfalfae* gene models and RNA-Seq analysis

Using a customized Genialis Platform (Genialis, Slovenia; <u>https://www.genialis.com/genialis-platform/</u>), the gene models of the *V. nonalfalfae* reference genome [22] were translated into putative proteins with the ExPASy Translate tool [100] and ORF Finder [101]. The general characteristics of putative proteins (molecular weight, number of amino acids (aa), percentage of cysteines and isoelectric point) were predicted by the ProtParam tool [100]. Functional annotation of the predicted proteins was performed with HMMER searches [102] against CAZy [103,104], Pfam [105], and Superfamily [106], as well as with BLAST searches [107] against NCBI, KOG [108], *MEROPS* [109] and PHI databases [62], followed by Blast2GO [110] and KEGG [111,112] analyses. The overrepresentation of GO terms in the *V. nonalfalfae in-silico* secretome compared to proteome was assessed using a hypergeometric distribution test (HYPGEOM.DIST function in Excel) with a p-value < 0.05 and FDR < 0.05.

RNA-sequencing of *V. nonalfalfae* mild and lethal pathotypes was performed by IGA Technology Service (Udine, Italy) using Illumina Genome Analyzer II. For this purpose, total RNA, enriched for the polyA mRNA fraction, was isolated in three biological replicates from fungal mycelia of mild and lethal strains grown in xylem-simulating media according to [98]. Illumina raw sequence reads were deposited at NCBI (Bioproject PRJNA283258). RNA-Seq analysis was performed using CLC Genomics Workbench tools (Qiagen, USA). Differentially expressed genes between lethal and mild fungal pathotype were identified as those with fold change FC ≥ 1.5 or FC ≥ – 1.5 (*p* ≤ 0.05; FDR ≤ 0.05).

From our previous RNA-Seq data of compatible and incompatible interactions between hop and *V. nonalfalfae* [45], fungal transcripts expressed at a least one time point (6, 12, 18 and 30 dpi) and one hop cultivar (susceptible ‘Celeia’ or resistant ‘Wye Target’) were filtered out and data were presented as a matrix of log_2_CPM (counts per million – number of reads mapped to a gene model per million reads mapped to the library) expression values. These genes were considered as expressed *in planta.*

### 5.3 Secretome prediction and comparison

The *V. nonalfalfae in silico* secretome was determined using a customized Genialis Platform (Genialis, Slovenia) according to the method described in [113], which reportedly gives 83.4% accuracy for fungal secreted proteins. It combines SignalP4.1 [114], WolfPsort [47] and Phobius [46] for N-terminal signal peptide prediction, TMHMM (<u>http://www.cbs.dtu.dk/services/TMHMM/</u>) for eliminating membrane proteins (allowing one transmembrane (TM) domain in the first 60 aa) and PS-Scan [115] for removing proteins with ER targeting sequence (Prosite: PS00014). In addition, we used LOCALIZER [49] to predict effector protein localization to chloroplasts and mitochondria, while proteins with a nuclear localization signal were determined with NucPred (likelihood score >0.80) [116] and PredictNLS [117]. Localization of effector proteins to the apoplast was predicted by ApoplastP [48] based on enrichment in small amino acids and cysteines, as well as depletion in glutamic acid.

To compare the composition of the *V. nonalfalfae* secretome to other closely related plant pathogenic *Verticillium* species, protein coding sequences of *Verticillium dahliae* JR2, *Verticillium longisporum* GCA_001268145 and *Verticillium alfalfae* VaMs.102 from the Ensembl Fungi database (<u>http://fungi.ensembl.org/info/website/ftp/index.html</u>) were used and secretome predictions, HMMER searches against CAZy database and blastp searches against the *MEROPS* database were performed using the same pipeline as for *V. nonalfalfae.* Two way ANNOVA followed by Tukey’s multiple comparisons test (p-value < 0.05) in GraphPad Prism 7.03 (GraphPad Software, Inc., USA) was used to find differences between sets of fungal proteins.

### 5.4 Refinement of *V. nonalfalfae* secretome and selection of CSEPs

A refinement of total *V. nonalfalfae in silico* secretome (Figure 1) was done to maintain only proteins, transcripts of which were expressed *in planta* according to the RNA-Seq analysis. Proteins with carbohydrate enzymatic activities (CAZy screening) were excluded from further analysis and additional filtering was applied based on the presence of PFAM domains. CSEPs were identified as proteins having known effector-specific domains [9,12,13,16,118], NLS signal or as proteins with no PFAM domains.

In the second step, the number of secreted proteins was narrowed down using the following criteria: proteins determined in the lethal pathotype-specific region identified by comparative genomics of mild and lethal *V. nonalfalfae* strains from three geographic regions [22]; proteins differentially expressed in lethal compared to mild *V. nonalfalfae* strains grown in xylem-simulating media as determined by RNA-Seq; *V. nonalfalfae* secreted proteins analyzed by 2D-DIGE and identified by MALDI-TOF/TOF MS [44]; proteins with sequence similarity to experimentally verified pathogenicity, virulence and effector genes in the PHI (Pathogen-host Interaction) database [62] and putative effector proteins predicted by EffectorP software [16].

### 5.5 Quantitative real time PCR and fungal biomass quantification

Susceptible ‘Celeia’ and resistant ‘Wye Target’ hop varieties were inoculated by the root dipping method [119] with *V. nonalfalfae* spores. Total RNA was isolated from the roots and stems of infected plants (6, 12 and 18 dpi) or mock-inoculated plants using a MagMAX total RNA isolation kit (Life Technologies, USA). The quality and quantity of RNA was assessed on an Agilent 2100 Bioanalyzer (Agilent Technologies, Germany). Total RNA was transcribed to cDNA with a High Capacity cDNA Reverse Transcription kit (Applied Biosystems, USA). Real-time PCR reactions were performed on 7500 Fast Real Time PCR Systems (Life Technologies, USA) using the FastStart SYBR Green master mix (Roche, Switzerland) and primers (Table S8) designed by Primer Express 3.0 software (Thermo Fisher Scientific, USA). For each of the 44 top priority CSEPs, gene expression was analyzed on pooled samples containing the roots of five individual plants, in two technical replicates. The highest expressed CSEPs were also analyzed in five biological and two technical replicates per sample group. The CSEPs’ gene expression was calculated by the comparative C_T_ method [120]. The cDNA from *V. nonalfalfae* mycelium cultivated on ½ CD was used as a reference sample. *V. nonalfalfae* DNA topoisomerase *(Vna Un148)* and splicing factor 3a2 *(Vna8.801)* were selected as the best endogenous control genes according to GeNorm analysis [121] and fungal biomass normalization. For the latter, fungal DNA was extracted from infected hop using CTAB and quantified by qPCR as described in [122]. The expression of control genes was compared to fungal biomass in infected hop using Pearson’s correlation coefficient.

### 5.6 Construction of knockout vectors and preparation of *V. nonalfalfae* knockout mutants

CSEPs knockout mutants were prepared according to Frandsen’s protocol [123]. The plasmid vector pRF-HU2 was first linearized with Nt.*Bbv*CI and *Pac*I (New England BioLabs, USA) and purified with Illustra GFX PCR DNA and Gel Band Purification Kit (GE Healthcare, UK). Homologous gene sequences were then amplified with PCR using a PfuTurbo Cx Hotstart DNA polymerase (Agilent Technologies, USA) and the following settings: 95°C for 2 minutes, 30 cycles: 95°C for 30s, 55°C for 30s, 72°C for 1 minute; 75°C for 10 minutes. PCR products were purified with Illustra GFX PCR DNA and a Gel Band Purification Kit (GE Healthcare, UK) and ligated to linearized vector pRF-HU2 with USER enzyme (New England BioLabs, USA). Vector constructs were multiplied in *E. coli* DH5α cells and isolated with a High Pure Plasmid Isolation Kit (Roche, Life Science, USA).

*V. nonalfalfae* knockout mutants were generated with *Agrobacterium tumefaciens* mediated transformation (ATMT) [124] using acetosyringone (Sigma Aldrich, USA). The knockout vector constructs were electroporated with Easyject Prime (EQUIBIO, UK) into electro-competent *Agrobacterium tumefaciens* (LBA4404) cells. Positive colonies with a correct construct orientation were verified by colony PCR. Co-culture of transformed *A. tumefaciens* and *V. nonalfalfae* was carried out on IMAS media as described in [99]. Colonies were transferred on a cellophane membrane (GE Healthcare, UK) to primary and secondary ½ CD selection medium with 150 mg/L timentin (Duchefa, The Netherlands) and 75 mg/L hygromycin (Duchefa, The Netherlands). Genomic DNA was isolated according to [125] from the remaining colonies and the knockout was verified with PCR. Transformed *V. nonalfalfae* conidia were stored in 30% glycerol at -80°C until testing.

### 5.7 Pathogenicity evaluation of *V. nonalfalfae* knockout mutants in hop

Before pathogenicity tests were carried out, fungal growth and conidiation were inspected as described previously [126]. Ten to fifteen plants of the Verticillium wilt susceptible hop cultivar ‘Celeia’ were inoculated at phenological stage BBCH 12 by 10-min root dipping in a conidia suspension of *V. nonalfalfae* knockout mutants as described previously [126]. Conidia of the wild type *V. nonalfalfae* lethal pathotype served as a positive control and sterile distilled water was used as a mock control. Verticillium wilting symptoms were assessed four to five times post inoculation using a disease severity index (DSI) with a 0-5 scale [42], and rAUDPC was calculated according to [63]. After symptom assessment, fungal re-isolation test and qPCR using *V. nonalfalfae* specific primers (Table S8) were performed to confirm infection of hops.

### 5.8 Statistics

The R package [127] was used for the statistical analysis of the pathogenicity assay of knockout mutants. Due to the different variability of rAUDPC values for the ‘isolate’ groups, a non-parametric approach was pursued. A Kruskal-Wallis test was used, followed by multiple comparison test with Bonferroni correction. To understand how the time post inoculation with *V. nonalfalfae* affects hop health, a simple logistic growth model [128] was fitted to DSI values for the groups under study.

## 6

### List of Abbreviations

AA –: proteins with auxiliary activities
ATMT –: *Agrobacterium tumefaciens* mediated transformation
CBM –: proteins with carbohydrate-binding modules
CE –: carbohydrate esterases
CSEPs –: candidate secreted effector proteins
CWDEs –: cell wall-degrading enzymes
DSI –: disease severity index
ETI –: effector triggered immunity
GH –: glycoside hydrolases
GT –: glycosyltransferases
KOG –: EuKaryotic Orthologous Groups
NLPs –: necrosis and ethylene-inducing protein (NEP-1)-like proteins
NLR –: nucleotide-binding leucine rich repeat
Nox –: NADPH oxidase complex
PAMPs –: pathogen-associated molecular patterns
PHI –: Pathogen-host interaction database
PL –: polysaccharide lyases
rAUDPC –: relative area under the disease progress curve
ROS –: reactive oxygen species
SSCPs –: small secreted cysteine-rich proteins
SSPs –: small secreted proteins
TM –: transmembrane domain
XSM –: xylem simulating medium

## 7 Declarations

### Ethics approval and consent to participate

Not applicable.

### Consent for publication

Not applicable.

### Availability of data and materials

All data generated or analyzed during this study are included in this published article and its supplementary information files. *Verticillium nonalfalfae* genomic data are available at NCBI under BioProject <u>PRJNA283258</u>, while transcriptome data of *Verticillium nonalfalfae* interaction with hop can be retrieved from NCBI Bioproject <u>PRJEB14243</u>.

### Authors’ Contributions

BJ conceived and coordinated the study and SB participated in its design. JJ performed *Verticillium nonalfalfae* genome assembly and gene model predictions. SR prepared the plant and fungal material and performed hop inoculations. KM and SB performed secretome analysis and RT-qPCR experiments. MF and KM prepared ATMT knockout mutants and performed the pathogenicity assays together with SR. KK performed statistical analysis of data. KM, SB and BJ analyzed and interpreted the data and drafted the manuscript. All authors read and approved the final manuscript.

## Acknowledgments

We express our sincere gratitude to Dr. Vasja Progar and the Genialis team for their help with bioinformatics. We acknowledge advice on data normalization in RT-qPCR experiments by Dr. Nataša Štajner from the University of Ljubljana. We thank Martin Cregeen for English language editing.

## 10 Additional files

<u>Additional file 1</u>. An Excel file containing 8 tables, each in a separate worksheet.

Table S1. *V. nonalfalfae* secretome predicted using Genialis bioinformatics platform. Gene IDs marked in bold represent 263 proteins in the CSEPs catalogue. 2D-DIGE, proteins secreted by mild and lethal strains of *V. nonalfalfae* growing in xylem simulating medium (XSM) were analysed by 2D-DIGE and identified by MALDI-TOF/TOF MS [44]; RNA-Seq, differential fungal gene expression (fold change, FC ≥ 1.5 or FC ≤ – 1.5) of *V. nonalfalfae* lethal versus mild fungal pathotype growing in XSM; LOCALIZER, a tool for subcellular localization prediction of both plant and effector proteins in the plant cell [49]; ApoplastP, a tool for prediction of effectors and plant proteins in the apoplast using machine learning [48].

Table S2. KOG functional annotation of *V. nonalfalfae* predicted secretome. KOG, a database of euKaryotic Orthologous Groups from NCBI that allows identification of ortholog and paralog proteins [108].

Table S3. *MEROPS* classification of predicted *V. nonalfalfae* secretome. *MEROPS,* a database of peptidases and the proteins that inhibit them [109].

Table S4. Assignment of KEGG accessions to the predicted *V. nonalfalfae* secreted proteins. KEGG, the Kyoto Encyclopedia of Genes and Genomes is a database resource for understanding high-level functions and utilities of the biological systems [111].

Table S5. List of PFAM domains identified in the predicted *V. nonalfalfae* secretome. PFAM, a collection of protein families, represented by multiple sequence alignments and hidden Markov models (HMMs) [105].

Table S6. List of *V. nonalfalfae* CSEPs, including EffectorP prediction and similarity to effectors in the PHI database. EffectorP, a machine learning prediction program for fungal effectors [16]; PHI, Pathogen-Host Interaction database, which contains expertly curated molecular and biological information on genes proven to affect the outcome of pathogen-host interactions [62].

Table S7. Bulk expression of best ranked *V. nonalfalfae* candidate effector genes in hop roots. Gene expression was calculated by the comparative C_T_ method [120]. Hop plants were inoculated by the root dipping method with *V. nonalfalfae* conidia and sampled at 6, 12 and 18 days post inoculation. Analyzed samples contained the roots of five individual plants from either susceptible hop variety ‘Celeia’ (CE) or resistant ‘Wye Target’ (WT). cDNA from *V. nonalfalfae* mycelium cultivated on ½ Czapek Dox (CD) medium was used as a reference sample. *V. nonalfalfae* DNA topoisomerase *(VnaUn.148)* and splicing factor 3a2 *(Vna8.801)* were used as endogenous control genes. Numbers represent log2 fold changes in the expression of genes in infected plants at indicated time points compared to gene expression in ½ CD medium. na, not available

Table S8. List of primers used in this study. ^a^, oligonucleotide designed according to template [123]; ^b^, oligonucleotide that amplifies the promoter region of the target gene; ^c^, oligonucleotide that amplifies the terminator region of the target gene; ^d^, oligonucleotide that amplifies the target gene for knockout (KO) selection; ^e^, oligonucleotide that amplifies genomic and vector sequences for KO selection; ^f^, *V. nonalfalfae* lethal pathotype-specific marker [97].

<u>Additional file 2</u>. A figure in PDF format. Enrichment of carbohydrate metabolic processes and peptidase activity in the predicted *V. nonalfalfae* secretome compared to proteome. GO terms for biological process (A) and molecular function (B) are presented at GO level 4. Relative abundance of gene ontology (GO) terms determined by Blast2GO is expressed in percentages of predicted fungal secretome and proteome, respectively.

<u>Additional file 3</u>. A figure in PDF format. A heatmap displaying the expression patterns of *V. nonalfalfae* genes during infection of susceptible ‘Celeia’ and resistant hop ‘Wye Target’. Fungal transcripts were first identified by mapping of reads with at least 90% sequence identity and 90% sequence coverage to the *V. nonalfalfae* reference genome [22] using CLC Workbench. Normalization by trimmed mean of M values (TMM) [129] was performed to eliminate composition biases between libraries. Read counts were converted into log2-counts-per-million (logCPM) values and a cutoff of CPM >1 was chosen. Color scale bar represents the logCPM values, with darker red color meaning higher expression values.

<u>Additional file 4</u>. A figure in PDF format. Some *V. nonalfalfae* CSEPs deletion mutants exhibit unaffected pathogenicity. Relative area under the disease progress curve (rAUDPC) is presented for the three *V. nonalfalfae* deletion mutants (each in four replicates) and the wild type fungus. Depicted are mean values ± SEM (n=10). Analysis of variance was first performed by Levene’s test, followed by Dunnett’s test to compare each treatment (knockout) with a single control (wild type); however, no statistical differences were found at α level of 5%.

